# Anionic lipids stabilize etioplast prolamellar bodies and facilitate thylakoid biogenesis during etioplast-to-chloroplast differentiation

**DOI:** 10.64898/2026.10.31.763292

**Authors:** Akiko Yoshihara, Risa Uwatoko, Keiko Kobayashi, Noriko Nagata, Koichi Kobayashi

## Abstract

Etioplasts of dark-grown angiosperm seedlings contain prolamellar bodies (PLBs), lattice-like membranes that are quickly transformed into thylakoids upon illumination. Phosphatidylglycerol (PG) and sulfoquinovosyldiacylglycerol (SQDG) are major anionic lipids in these membranes, but their roles in etioplast-to-chloroplast differentiation remain unclear. We analyzed *Arabidopsis thaliana* mutants with reduced PG biosynthesis (*pgp1-1*), complete SQDG deficiency *(sqd1*), and the *pgp1-1 sqd1* double mutant. Although *pgp1-1* etioplasts showed a loosened PLB lattice when fixed under white light, they retained regular, wild-type-like lattices when fixed under infrared light, suggesting that their PLBs were more susceptible to light-induced disassembly. The rapid PLB disassembly in *pgp1-1* upon illumination was accompanied by rapid conversion of dark-accumulated protochlorophyllide to chlorophyll and lamellar membrane formation, suggesting a link between membrane remodeling and chlorophyll formation. However, subsequent chlorophyll accumulation through de novo biosynthesis was retarded in illuminated *pgp1-1* seedlings, whereas photosynthetic electron transport was quickly activated. Although *sqd1* showed no obvious defects, *pgp1-1 sqd1* exhibited more severe defects than *pgp1-1*, particularly in internal membrane formation, chlorophyll accumulation, photosynthetic activation, and photosynthetic gene expression. These findings suggest that PG is crucial for membrane-associated processes during etioplast-to-chloroplast differentiation, while SQDG may help maintain chloroplast development when PG availability is limited.

## Introduction

Cotyledon cells of dark-grown angiosperm seedlings develop chloroplast precursors called etioplasts. Etioplasts contain a three-dimensional lattice of branched membrane tubules termed prolamellar bodies (PLBs). Protochlorophyllide (Pchlide), an intermediate of chlorophyll (Chl) biosynthesis, accumulates in PLBs instead of Chl. Most Pchlide is bound at the catalytic site of light-dependent NADPH-Pchlide oxidoreductase (LPOR), which also functions as a membrane-remodeling protein essential for PLB formation (Schoefs and Franck, 2003; Solymosi and Schoefs, 2010; Nguyen *et al*., 2021). LPOR forms oligomeric ternary complexes with Pchlide and NADPH, and these complexes helically coat the PLB membranes (Floris and Kühlbrandt, 2021).

Upon illumination, Pchlide is instantaneously converted into chlorophyllide (Chlide) by LPOR, followed by the gradual degradation of LPOR, leading to the disassembly of PLBs into lamellar membranes (Solymosi and Schoefs, 2010; Kowalewska *et al*., 2016). Although Chl synthase, which catalyzes the esterification of Chlide with a phytol chain, is considered inactive when the PLB structure is highly organized, it becomes activated upon PLB disassembly (Lindsten *et al*., 1990). In parallel, illumination upregulates the expression of photosynthesis-associated genes, thereby promoting Chl and photosynthetic protein biosynthesis (Ma *et al*., 2001; Pipitone *et al*., 2021). Along with LPOR degradation, the accumulation of photosynthetic membrane protein complexes such as light-harvesting complex II (LHCII) drives the transformation of the tubular PLB lattice into stacked thylakoid membranes and the differentiation of etioplasts to chloroplasts (Fujii *et al*., 2019*b*).

The composition of pigments and proteins substantially changes during the PLB-to-thylakoid transformation (Blomqvist *et al*., 2008; Plöscher *et al*., 2011; Garmash *et al*., 2013; Pipitone *et al*., 2021). By contrast, the lipid composition of PLBs is similar to that of thylakoid membranes (Selstam and Sandelius, 1984; Dorne *et al*., 1990), and thus PLBs are regarded as a lipid reservoir for the formation of initial thylakoid membranes (Solymosi and Schoefs, 2010; Fujii *et al*., 2019*b*). The PLB lipid bilayer is mainly composed of two galactolipids, monogalactosyldiacylglycerol (MGDG) and digalactosyldiacylglycerol (DGDG), which account for ∼50% and ∼30% of total PLB lipids, respectively (Selstam and Sandelius, 1984). These galactolipids are required for etioplast development (Fujii *et al*., 2017, 2018) and for etioplast-to-chloroplast differentiation (Fujii *et al*., 2019*a*). In addition, PLBs contain sulfoquinovosyldiacylglycerol (SQDG), a sulfur-containing glycolipid, and the phospholipid phosphatidylglycerol (PG), each accounting for ∼10 mol% of total PLB lipids (Selstam and Sandelius, 1984).

PG and SQDG are anionic lipids whose polar head groups are negatively charged at physiological pH. PG is indispensable for photosynthesis in oxygenic photosynthetic organisms including cyanobacteria, algae, and plants (Wada and Murata, 2007; Kobayashi *et al*., 2024). Many PG molecules are specifically associated with photosystem I (PSI) and photosystem II (PSII), where they likely play essential roles in the structure and function of these complexes (Yoshihara and Kobayashi, 2022). Accordingly, in *Arabidopsis thaliana*, a T-DNA insertional knockout mutation (*pgp1-2*) of phosphatidylglycerophosphate synthase 1 (PGP1), which results in the complete loss of plastidic PG biosynthesis, causes severe impairments of photosynthetic reactions and thylakoid membrane development (Hagio *et al*., 2002; Kobayashi *et al*., 2016). Meanwhile, an approximately 80% reduction in PGP1 activity caused by a single amino acid substitution (*pgp1-1*) results in a ∼30% decrease in leaf PG content and a pale-green phenotype, without a severe impairment of photosynthetic capacity (Xu *et al*., 2002). However, additional loss of SQDG biosynthesis in the *pgp1-1* background leads to defects in photosynthetic electron transport, Chl accumulation, thylakoid development, and growth under light conditions (Yu and Benning, 2003; Yoshihara *et al*., 2021). Because the loss of SQDG biosynthesis alone does not impair these processes in *A. thaliana* (Yu *et al*., 2002; Yoshihara *et al*., 2021), SQDG likely compensates for some functions of PG when PG levels are insufficient. Indeed, SQDG content increases as PG content decreases under phosphate-deficient conditions, thereby maintaining the total level of anionic lipids (Essigmann *et al*., 1998). Because mutants lacking SQDG biosynthesis exhibit decreased photosynthetic activity under phosphate-deficient conditions (Yoshihara *et al*., 2021), the compensatory role of SQDG may be particularly important during phosphate deficiency.

We have recently reported that PG also plays an important role in etioplast development, whereas SQDG supports the function of PG (Yoshihara *et al*., 2024). Etioplasts in *pgp1-1* cotyledons grown in the dark show a loosened PLB lattice structure, impaired Pchlide biosynthesis particularly at the Mg-insertion step, and altered behavior of Pchlide-LPOR-NADPH complexes. In contrast, the *sqd1* and *sqd2* mutants, which completely lack SQDG biosynthesis due to knockout mutations in *SQD1* and *SQD2*, respectively, show no obvious defects in etioplast development. However, the *pgp1-1 sqd1* and *pgp1-1 sqd2* double mutants exhibit more severe defects than *pgp1-1* during etioplast development. These findings suggest that anionic lipids, particularly PG, are required for membrane-protein organization even before the accumulation of photosynthetic membrane proteins in the dark (Yoshihara *et al*., 2024).

A recent review proposed a model in which lipid biosynthesis triggers Chl and photosynthetic protein biosynthesis and promotes thylakoid membrane biogenesis and photosynthetic activation (Fujii *et al*., 2025). During etioplast-to-chloroplast differentiation, galactolipid biosynthesis is required for the efficient organization of the Chl biosynthesis pathway and for photosynthetic gene expression in plastids and the nucleus (Fujii *et al*., 2019*a*, 2024). In contrast, it remains unclear whether anionic lipids play specific roles in these processes. In this study, we investigated the de-etiolation processes of dark-grown seedlings of *pgp1-1*, *sqd1*, and the *pgp1-1 sqd1*double mutant, including rapid conversion of Pchlide to Chl via Chlide, PLB disassembly, de novo Chl biosynthesis, photosynthetic activation, and photosynthetic gene expression. The results highlight the specific roles of plastid anionic lipids, particularly PG, in etioplast-to-chloroplast differentiation.

## Materials and methods

### Plant materials and growth conditions

The wild type Col-0 (WT), *sqd1* (Okazaki *et al*., 2009), *pgp1-1* (Xu *et al*., 2002), and *pgp1-1 sqd1* plants (Yoshihara *et al*., 2021), all in the Columbia background of *A. thaliana*, were used in this study.

Surface-sterilized seeds were sown on half-strength Murashige and Skoog medium (pH 5.7 adjusted with KOH) containing 1% (w/v) sucrose and solidified with 0.8% (w/v) agar, cold-treated at 4 °C for 3–4 d in the dark, and then exposed to white light (∼80 μmol photons m ² s ¹) at 23 °C for 4 h to synchronize germination. Seedlings grown in the dark for 4 d were either used for experiments or subsequently illuminated with continuous white light (∼80 μmol photons m ² s ¹) at 23 °C for deetiolation.

### Transmission electron microscopy and scanning electron microscopy analysis

The samples were processed for electron microscopy based on the procedure reported by Fujii *et al*. (2017), with minor modifications. Except for samples fixed under infrared light, cotyledons were dissected immediately after sampling and immersed in a fixative containing 4% glutaraldehyde and 4% paraformaldehyde at 4°C overnight. After fixation, the samples were postfixed in 2% OsO_4_ supplemented with 1.5% K_3_[Fe(CN)_6_] at 4°C for 2 h, dehydrated through an ethanol series, transferred to propylene oxide, infiltrated with Spurr’s resin, and embedded.

For fixation under infrared light, dark-grown seedlings were handled under 850-nm infrared illumination (GR10-IR, SHODENSHA, Osaka, Japan) using an infrared camera (GR200HD2-IR, SHODENSHA). Because fine dissection was difficult under these conditions, whole seedlings were first immersed in the same fixative, wrapped in aluminum foil, and stored at 4°C for 2 days to allow fixation without exposure to visible light. The fixed seedlings were then dissected and processed for postfixation, dehydration, resin infiltration, and embedding as described above.

Ultrathin sections (80 nm thick) were stained with uranyl acetate and lead citrate and observed by transmission electron microscopy (TEM; JEM-1400, JEOL, Tokyo, Japan) at high magnification to examine fine structural details of plastid internal membranes. For scanning electron microscopy (SEM), which was used to examine larger tissue areas containing numerous plastid profiles, the stained sections were further coated with OsO_4_ and observed by field-emission SEM (SU8220, Hitachi, Tokyo, Japan). Plastid profiles in the SEM images were manually classified and quantified based on their internal membrane organization. Plastid profiles in which internal membranes were absent or could not be clearly distinguished were excluded from the analysis. In addition, to assess the three-dimensional organization of *pgp1-1 sqd1* plastids, serial ultrathin sections, each 80 nm thick, were prepared and observed by SEM.

### In situ low-temperature fluorescence spectroscopy

For analysis of the time-dependent changes in low-temperature (77 K) fluorescence spectra during dark incubation after a single flash, approximately 30 etiolated seedlings grown for 4 d in the dark were placed in a clear microtube under dim green light, exposed to a 0.7-ms single flash (PZ42X, Sunpak), and kept in darkness at 13 °C for the indicated time. The microtube was then placed in a liquid nitrogen-filled Dewar flask positioned in the cuvette holder of a fluorometer (RF-6000, Shimadzu, Kyoto, Japan). Fluorescence emission spectra were recorded in liquid nitrogen under 440-nm excitation.

For analysis of light-induced transitions of low-temperature fluorescence spectra, approximately 30 etiolated seedlings were placed in a clear microtube, exposed to continuous white light (∼80 μmol photons m ² s ¹) at 23 °C for the indicated time after a flash, and used for recording fluorescence emission spectra under 435-nm excitation as described above.

### Immunoblot analysis

Seedlings grown for 4 d in the dark were exposed to white light (∼80 μmol photons m^−2^ s^−1^) for 5 minutes and then frozen in liquid nitrogen. Immunoblot analysis, from total protein extraction to detection and quantification of total LPOR proteins, was performed as described previously (Yoshihara *et al*., 2024).

### Determination of porphyrin content

To determine the ratio of Chl *a* to Chlide *a* after a flash, 4-d-old etiolated seedlings were incubated in the dark for the indicated time after a single flash, immediately frozen in liquid nitrogen, homogenized in N,N-dimethylformamide, and analyzed as described previously (Fujii *et al*., 2017) using the high performance liquid chromatography (HPLC) system (Yoshihara *et al*., 2024). The ratio of Chl *a* to Chlide *a* was determined from respective fluorescence peak areas measured at 670 nm under 430-nm excitation.

To determine the porphyrin composition after continuous illumination, 4-d-old etiolated seedlings were exposed to white light (∼80 μmol photons m^−2^ s^−1^) for the indicated time, extracted in N,N-dimethylformamide at 4 °C in the dark for 1 d, and analyzed by HPLC. Pchlide content was determined by measuring the fluorescence at 634 nm under 440 nm excitation and using a known concentration of Pchlide as a standard. To determine Chlide *a* and Chl *a* contents, peak areas in the fluorescence chromatograms recorded at 670 nm under 430-nm excitation were converted to amounts using a calibration based on the photoactive Pchlide content determined before illumination, which corresponds to the Chl(ide) *a* content formed after the flash.

To analyze Chl accumulation, 4-d-old etiolated seedlings were exposed to white light (∼80 μmol photons m^−2^ s^−1^) for the indicated time and then incubated in 80% (v/v) acetone at 4 °C in the dark for 1 d. Chl content was determined by measuring fluorescence emission at 670 nm under 435-nm excitation using a spectrofluorometer (RF-6000), with a known concentration of Chl as a standard. At 0 h, the amount of Pchlide described in Yoshihara *et al*. (2024) was used instead of Chl content. The Chl accumulation rate was calculated by dividing the Chl content accumulated during the indicated time period by the duration of the period.

### Photosynthetic efficiency analysis

Chl fluorescence parameters were analyzed using an imaging pulse amplitude modulation (PAM) fluorometer (IMAGING-PAM MINI, Walz, Effeltrich, Germany) and ImagingWinGigE software. Etiolated seedlings exposed to continuous white light (∼80 μmol photons m^−2^ s^−1^) for the indicated time were dark-adapted for 5 min. Minimal fluorescence (*F_o_*) was determined using the lowest measuring light (460 nm, ∼0.2 μmol photons m^−2^ s^−1^ at a frequency of 2 Hz), and maximal fluorescence (*F_m_*) was obtained by applying a saturating pulse (460 nm, ∼3400 μmol photons m^−2^ s^−1^ for 720 ms). The seedlings were then illuminated with actinic light (460 nm, ∼75 μmol photons m^−2^ s^−1^) for 10 min, and a second saturating pulse was applied to determine the steady-state fluorescence (*F_s_*) and maximal fluorescence under actinic illumination *(F*′*_m_*), respectively. The maximum quantum yield of PSII (*F_v_*/*F_m_*) was calculated as (*F_m_* − *F_o_*)/*F_m_*. The effective quantum yield of PSII (Φ*_PSII_*) and the quantum yields of regulated (Φ*_NPQ_*) and non-regulated energy dissipation (Φ*_c_*) were calculated according to Guadagno *et al*. (2010) as follows: Φ*_PSII_* = (*F*′*_m_* − *F_s_*)/*F*′*_m_*, Φ*_NPQ_* = *F_s_*/*F*′*_m_* − *F_s_*/*F_m_*, and Φ*_c_* = *F_s_*/*F_m_*.

### Reverse transcription quantitative PCR analysis

mRNA level was quantified as described (Yoshihara et al., 2024) with some modifications. Complementary DNA was amplified by 2-step thermal cycling consisting of an initial denaturation step at 95 °C for 30 s followed by 40 cycles of 5 s at 95 °C and 30 s at 60 °C with the use of the THUNDERBIRD Next SYBR qPCR Mix (TOYOBO, Osaka, Japan) and 200 nM gene-specific primers (Supplementary Table S1). The thermal cycling and signal detection were performed in duplicate using the CFX Connect (Bio-Rad, Hercules, CA, USA). The relative abundance of all transcripts amplified was normalized to the mean expression levels of the reference genes *ACTIN8* and *UBIQUITIN11* according to Pfaffl (2001).

### Statistical analysis

For multiple comparison tests, one-way ANOVA with Tukey’s honestly significant difference post hoc test was performed using the multcomp R package (Hothorn *et al*., 2008). A significance level of P < 0.05 was used.

### Accession numbers

Sequence data of the genes investigated in this article can be found in The Arabidopsis Information Resource under the following accession numbers: *PGP1* (AT2G39290), *SQD1* (AT4G33030), *ACT8* (AT1G49240), *UBQ11* (AT4G05050), *CHLH* (AT5G13630), *HEMA1* (AT1G58290), *LHCA4* (AT3G47470), *LHCB6* (AT1G15820), *psaA* (ATCG00350), and *psbA* (ATCG00020).

## Results

### PLBs deficient in anionic lipids are prone to light-induced disassembly

We previously reported that cotyledon cells of dark-grown *pgp1-1* seedlings contained loosened PLBs in etioplasts (Yoshihara *et al*., 2024), resembling those observed in WT etioplasts after ∼1 h of illumination (Kowalewska *et al*., 2016). In the previous study, etiolated cotyledons were fixed under white light, which took around 5 min (Yoshihara *et al*., 2024). We hypothesized that the short-term exposure to white light during fixation caused rapid PLB disassembly in *pgp1-1* but not in WT. To test this hypothesis, we fixed etiolated seedlings under infrared light (850 nm), at a wavelength well beyond the absorption bands of Pchlide–LPOR complexes (Shibata, 1957).

In dark-grown WT and *sqd1* seedlings fixed under infrared light, etioplasts exhibited highly regular PLB lattices similar to those observed under white-light fixation (Fig. 1A). Even in *pgp1-1* seedlings, most etioplasts showed regular lattice structures of PLBs under infrared fixation, in contrast to their loosened appearance under white light (Yoshihara *et al*., 2024). These data suggest that *pgp1-1* forms WT-like PLBs in darkness, but that their lattice structures are rapidly disassembled upon brief light exposure during fixation. Nevertheless, quantitative classification of internal membrane structures revealed that 5.4% (13 of 242) etioplast profiles in ultrathin sections of *pgp1-1* exhibited disordered internal membrane networks instead of lattice structures (Fig. 1B), suggesting that decreased PG biosynthesis results in heterogeneous defects in PLB lattice formation among etioplasts.

**Figure 1.**
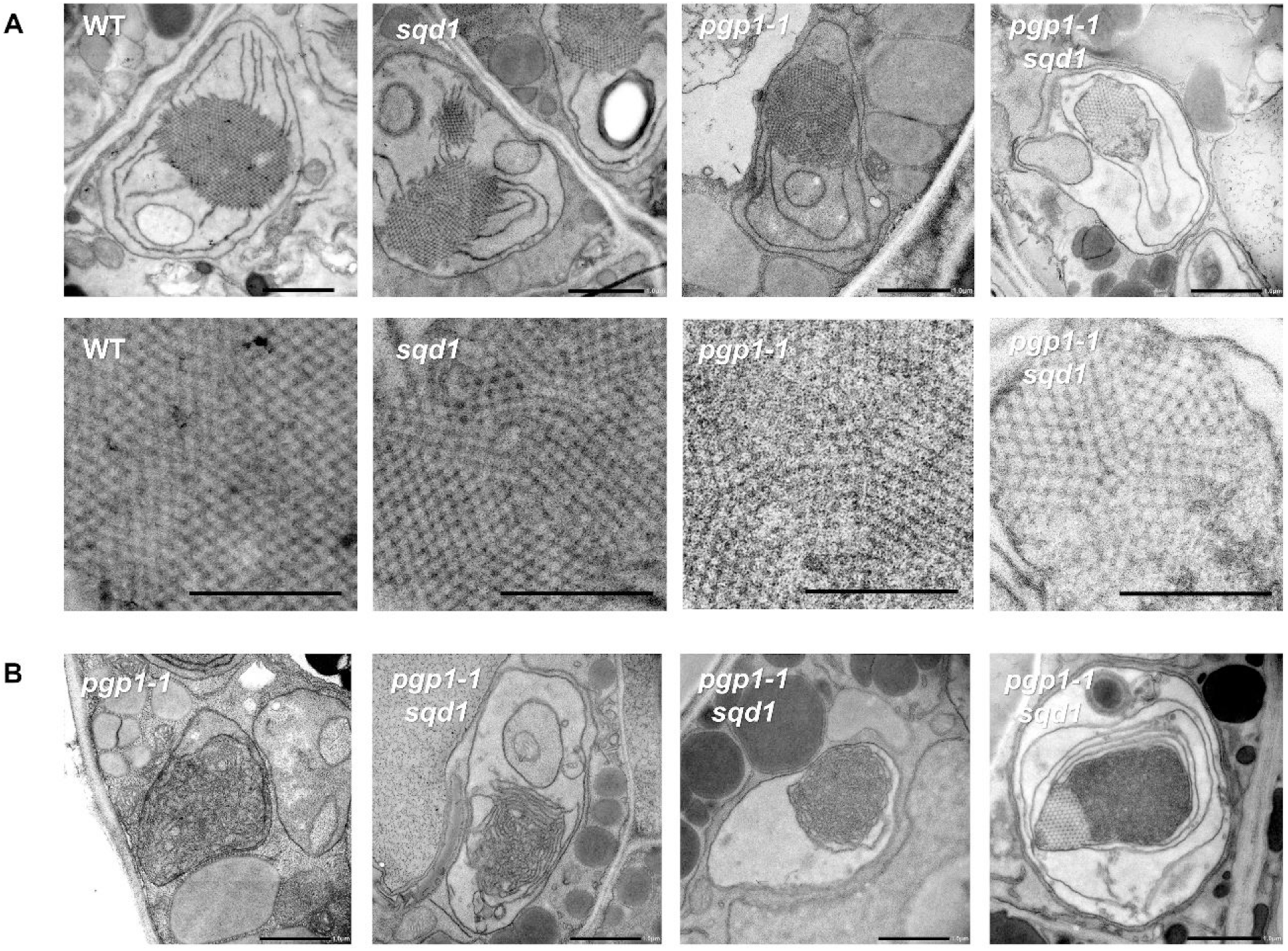
Membrane structures of etioplasts in 4-d-old etiolated cotyledons of wild-type (WT) and anionic lipid mutants. **(A)** Ultrastructures of etioplasts (upper panel) and the enlarged views of prolamellar body (PLB) lattices (lower panel) in cotyledons of dark-grown WT, *sqd1*, *pgp1-1*, and *pgp1-1 sqd1* seedlings fixed under infrared light. Bars = 1.0 μm (upper panel) and 0.5 μm (lower panel). In *pgp1-1 sqd1*, only 2.1% of etioplast profiles in ultrathin sections exhibited regular PLB lattices throughout the structure. **(B)** Ultrastructures of etioplasts containing disordered membrane networks in cotyledons of *pgp1-1* and *pgp1-1 sqd1* seedlings fixed under infrared light. Disordered internal membrane networks, rather than regular lattice structures, were observed in 5.4% of *pgp1-1* and 94.6% of *pgp1-1 sqd1* etioplasts in ultrathin sections. Bars = 1.0 μm. All images are high-resolution montages constructed from multiple images.

Our previous study also showed that etioplasts in the *pgp1-1 sqd1* double mutant fixed under white light exhibited disordered membrane networks or dense membrane aggregates instead of well-organized PLB lattices (Yoshihara *et al*., 2024). When fixed under infrared light, 94.8% (183 of 193) etioplast profiles in ultrathin sections of *pgp1-1 sqd1* exhibited only irregular membrane structures lacking PLB lattices, whereas 3.1% (6 of 193) showed irregular membranes containing partial lattice structures and 2.1% (4 of 193) exhibited regular PLB lattices throughout the structure (Fig. 1). However, three-dimensional analysis based on serial sections revealed a higher proportion of lattice-containing structures: 22.1% (43 of 194) of etioplasts contained partial lattice structures, whereas 5.6% (11 of 194) exhibited only regular lattices. Serial section analysis further showed that the partial lattice structures were not formed as a central core enclosed by irregular membranes, but instead occupied part of the internal membrane system, extending from the periphery to the interior (Supplementary Video S1). Considering that no lattice structures were observed in *pgp1-1 sqd1* etioplasts when fixed under white light (Yoshihara *et al*., 2024), these lattices are likely disassembled rapidly during the fixation process, as in *pgp1-1*.

### Anionic lipid deficiency accelerates dissociation of oligomeric Chlide-LPOR complexes and Chlide esterification

We previously reported that the peak shift of Chlide fluorescence following flash-induced Pchlide-to-Chlide conversion occurred more rapidly in *pgp1-1* and *pgp1-1 sqd1* than in WT (Yoshihara *et al*., 2024), suggesting a relationship between structural changes in PLBs and the dissociation of oligomeric Chlide-LPOR-NADP^+^ complexes (Franck *et al*., 1999; Schoefs and Franck, 2008). To examine how anionic lipids affect the initial phase of this dissociation process, etiolated seedlings of WT and anionic lipid mutants were kept in the dark at a reduced temperature (13 °C) for 1, 30, 60, or 120 s after a flash, and their fluorescence spectra were measured at 77 K.

In all samples, a fluorescence peak at ∼690 nm, originating from the Chlide-LPOR-NADP^+^ oligomer (Schoefs, 2001), appeared 1 s after a flash (Fig. 2A), indicating rapid photoconversion of Pchlide in the oligomeric ternary complex to Chlide, as observed previously (Yoshihara *et al*., 2024). In WT and *sqd1*, the 690-nm peak was red-shifted to ∼695 nm after 30 s (Fig. 2B), which presumably corresponds to fluorescence from the oligomeric Chlide-LPOR-NADPH complexes (El Hamouri *et al*., 1981; Oliver and Griffiths, 1982). In *pgp1-1* and *pgp1-1 sqd1*, the peak wavelengths after 30 s appeared to be shorter than those in WT and *sqd1*, but the differences were not significant. In WT, the peak had blue-shifted to 693.5 nm by 120 s, consistent with gradual dissociation of the oligomeric complexes. Similar peak shifts were observed in the anionic lipid mutants. However, the peak wavelengths in *pgp1-1* and particularly *pgp1-1 sqd1* were significantly shorter than that in WT, whereas *sqd1* showed a trend toward a longer peak wavelength than WT. In addition, a shoulder band at around 675 nm, which may originate from free Chlide released from the LPOR active site (Franck *et al*., 1999; Schoefs and Franck, 2008), appeared in all samples 30 s after illumination (Fig. 2A). The band was more intense in *pgp1-1* and even more pronounced in *pgp1-1 sqd1* than in WT and *sqd1* (Fig. 2C). In parallel with these band shifts, a small shoulder or peak was observed at ∼653 nm after 60 s (Fig. 2A), indicating the regeneration of photoactive Pchlide-LPOR complexes (Franck *et al*., 1999; Schoefs, 2001). The band was more distinct in *pgp1-1* and *pgp1-1 sqd1* than in WT and *sqd1*. These data suggest that the dissociation or structural rearrangement of Chlide-LPOR-NADPH complexes, release of free Chlide, and regeneration of photoactive Pchlide-LPOR complexes after a flash are accelerated in *pgp1-1* and particularly *pgp1-1 sqd1*.

**Figure 2.**
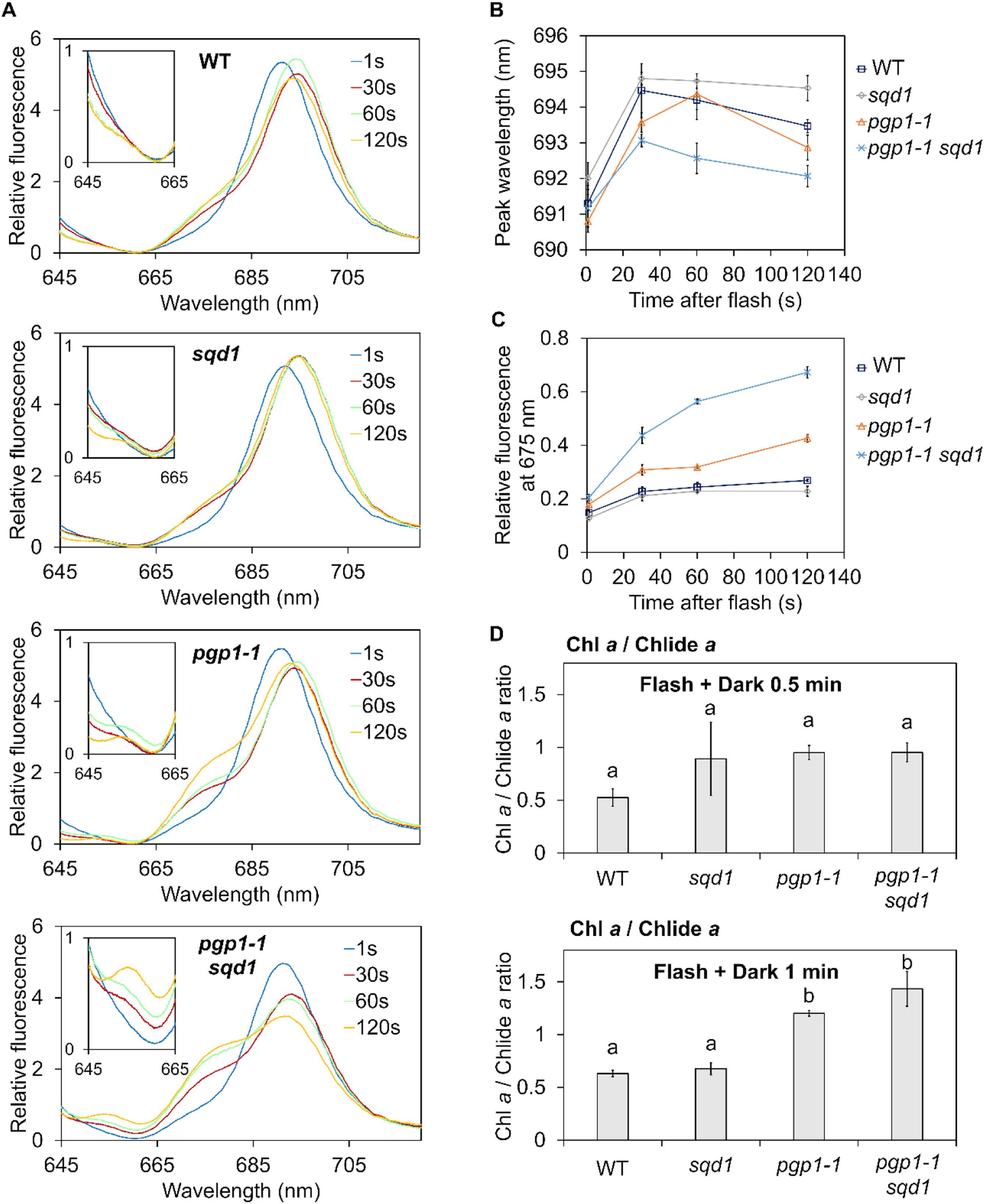
Rearrangement of oligomeric LPOR-pigment complexes and pigment metabolism in darkness following a flash in 4-d-old etiolated seedlings of wild type (WT) and anionic lipid mutants. (A) *In situ* low-temperature (77 K) fluorescence spectra of cotyledons after 1, 30, 60, and 120 s of dark incubation at 13 following a single flash. Fluorescence intensities were normalized to the mean fluorescence intensity over the 600–800 nm range. The insets show enlarged views of the spectra from 645 to 665 nm. Data are means from 3 independent experiments. **(B)** Changes in the fluorescence peak wavelength in the 690–696 nm region and **(C)** relative fluorescence intensity at 675 nm, normalized to the peak fluorescence intensity around 690 nm, during dark incubation. Data are means ± SE from 3 independent experiments. Statistical significance was assessed using one-way ANOVA followed by Tukey’s honestly significant difference (HSD) test. The results of the post hoc tests are provided in Supplementary Table S2. **(D)** Chl *a*/Chlide *a* ratio in etiolated seedlings after 0.5 and 1 min of dark incubation following a single flash. Data are means ± SE from 3 independent experiments. Different lowercase letters indicate significant differences among genotypes at each time point (p < 0.05, one-way ANOVA followed by Tukey’s HSD test).

Structural states of PLBs have also been reported to influence the activity of Chl synthase in esterifying Chlide with a phytyl chain (Lindsten *et al*., 1990, 1993). To evaluate Chlide esterification, we determined the Chl *a*/Chlide *a* ratio in etiolated seedlings after 0.5 and 1 min of dark incubation following the flash (Fig. 2D). At 1 min after the flash, *pgp1-1* and *pgp1-1 sqd1* showed more than a twofold higher Chl *a*/Chlide *a* ratio than WT and *sqd1*, suggesting accelerated phytylation of Chlide *a* to Chl *a* in these mutants.

### Anionic lipid deficiency causes rapid PLB-to-lamellar membrane transformation upon illumination

To investigate whether plastid anionic lipid deficiency affects PLB-to-thylakoid membrane transformation during etioplast-to-chloroplast differentiation, we performed time-course observations of plastid morphology upon illumination (Fig. 3A). In WT and *sqd1*, lamellar membranes gradually extended from the periphery of loosened PLBs, accompanied by progressive reduction in the size of PLB remnants over 6 h of illumination. While PLB remnants were still present in some plastids in WT and *sqd1* after 6 h, almost all PLB lattices had already transformed into lamellar membranes by 3 h in *pgp1-1*. After 9 h of illumination, no marked differences in the morphology of developing thylakoids were observed among WT, *sqd1*, and *pgp1-1*. These observations suggest that reduced PG biosynthesis accelerates PLB-to-lamellar membrane transformation but does not markedly affect subsequent thylakoid development, whereas loss of SQDG biosynthesis has little effect on these processes. In *pgp1-1*, disordered internal membrane networks and dense membrane aggregates, similar to those observed in dark-grown etioplasts, were also detected during illumination. Even after 24 h of illumination, 2.0% (6 of 296) of plastid profiles in ultrathin sections of *pgp1-1* cotyledons exhibited these abnormal membrane structures without forming thylakoids (Fig. 3B).

**Figure 3.**
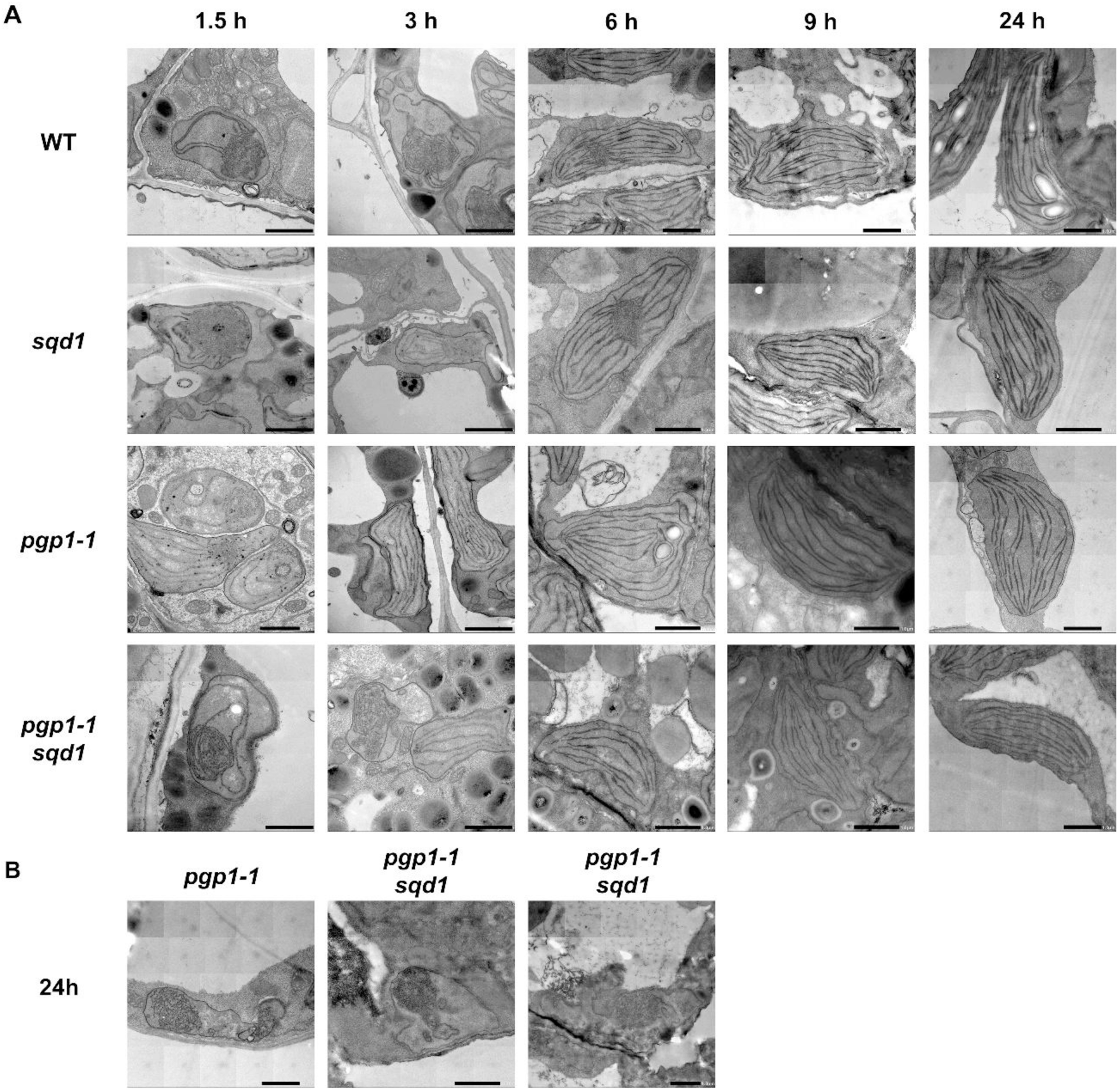
Time-course observation of plastid morphology in 4-d-old etiolated cotyledons of wild-type (WT) and anionic lipid mutants after illumination. **(A)** Ultrastructures of cotyledon plastids after 1.5, 3, 6, 9, and 24 h of continuous illumination in WT, *sqd1*, *pgp1-1*, and *pgp1-1 sqd1*. **(B)** Ultrastructures of plastids containing disordered membrane structures in *pgp1-1* and *pgp1-1 sqd1* after 24 h of illumination. Approximately 2% and 27% of plastid profiles in ultrathin sections of *pgp1-1* and *pgp1-1 sqd1*, respectively, exhibited abnormal membrane structures without forming thylakoid membranes. All images are high-resolution montages constructed from multiple images. Bars = 1.0 μm.

Rapid transformation from PLBs to lamellar membranes was also observed in *pgp1-1 sqd1*, with extended lamellar membranes and no detectable PLB remnants as early as 3 h after illumination (Fig. 3A). However, after 24 h of illumination, 26.8% (364 of 1358) of plastid profiles in ultrathin sections of *pgp1-1 sqd1* exhibited abnormal membrane structures instead of thylakoid membranes (Fig. 3B), suggesting that a subset of plastids retained abnormal membrane structures rather than developing normal thylakoid membranes, as observed in a few plastids in *pgp1-1*.

### Anionic lipid deficiency accelerates Pchlide-to-Chl conversion upon illumination

LPOR is gradually degraded upon illumination (Kay and Griffiths, 1983; Armstrong *et al*., 1995), and this degradation is thought to be associated with PLB disassembly (Fujii *et al*., 2019*b*). To assess whether the abnormally rapid PLB disassembly in *pgp1-1* and *pgp1-1 sqd1* results from accelerated LPOR degradation, we determined total LPOR levels after 5 min of illumination (Fig. 4A). LPOR content in *pgp1-1 sqd1* was 40% lower than in WT, whereas that in *sqd1* and *pgp1-1* was comparable to WT. A similar 40% reduction was previously observed in dark-grown *pgp1-1 sqd1* seedlings without illumination (Yoshihara *et al*., 2024), indicating that light-induced LPOR degradation is not notably accelerated in these mutants.

**Figure 4.**
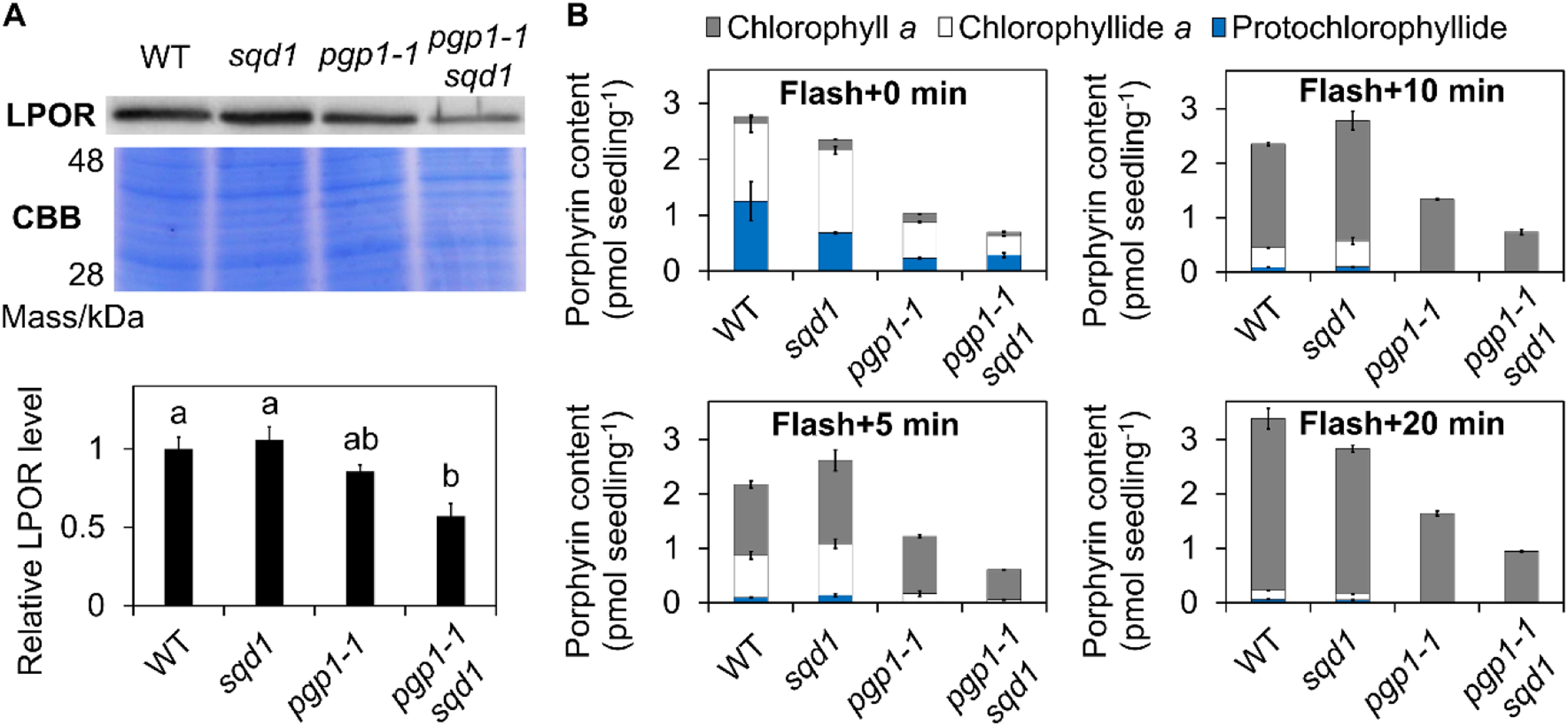
LPOR levels and porphyrin content in 4-d-old etiolated cotyledons of wild-type (WT) and anionic lipid mutants after illumination. **(A)** Immunodetection of total LPOR proteins (∼37 kDa), with Coomassie brilliant blue (CBB) staining shown as a loading control, and quantification of LPOR protein levels relative to the WT level in etiolated seedlings after 5 min of continuous illumination. Data are means ± SE from 3 independent experiments. Different lowercase letters indicate significant differences among genotypes (p < 0.05, one-way ANOVA followed by Tukey’s honestly significant difference test). **(B)** Chlorophyll *a*, chlorophyllide *a*, and protochlorophyllide content in 4-d-old etiolated seedlings after 0, 5, 10, and 20 min of continuous illumination following a single flash. Data are means ± SE from 3 independent experiments.

In etioplasts, Pchlide exists in photoactive and nonphotoactive forms (Schoefs, 2001). Immediately after a flash, both Chlide (derived from photoactive Pchlide) and Pchlide (nonphotoactive forms) were detected, with a small fraction of Chlide further converted to Chl (Fig. 4B). Total porphyrin levels after the flash were lower in *pgp1-1* and *pgp1-1 sqd1* than in WT and *sqd1*, consistent with reduced accumulation of Pchlide in these mutants during dark growth (Yoshihara *et al*., 2024). After 5 min of illumination following the flash, most porphyrin intermediates were converted to Chl in *pgp1-1* and *pgp1-1 sqd1*, with no detectable Pchlide remaining, whereas substantial and minor amounts of Chlide and Pchlide, respectively, remained in WT and *sqd1*. After 10 min, neither Pchlide nor Chlide was detected in *pgp1-1* and *pgp1-1 sqd1*, although small amounts of these intermediates were still detected in WT and *sqd1* even after 20 min. These results indicate that Pchlide-to-Chl conversion via Chlide occurs more rapidly in *pgp1-1* and *pgp1-1 sqd1*.

### Rapid PLB transformation in anionic lipid mutants does not enhance chloroplast development

To assess whether accelerated PLB-to-lamellar membrane transformation in *pgp1-1* and *pgp1-1 sqd1* promotes the establishment of chloroplast functions, we performed time-course analyses of Chl accumulation and photosynthetic capacity during deetiolation (Fig. 5). In *pgp1-1*, the rate of Chl accumulation was lower than those in WT and *sqd1*, particularly during the early stage of deetiolation, resulting in consistently lower Chl content throughout the illumination period. Furthermore, the *pgp1-1 sqd1* double mutant showed a markedly reduced rate of Chl accumulation and consequently a substantially lower Chl content after 24 h of illumination.

**Figure 5.**
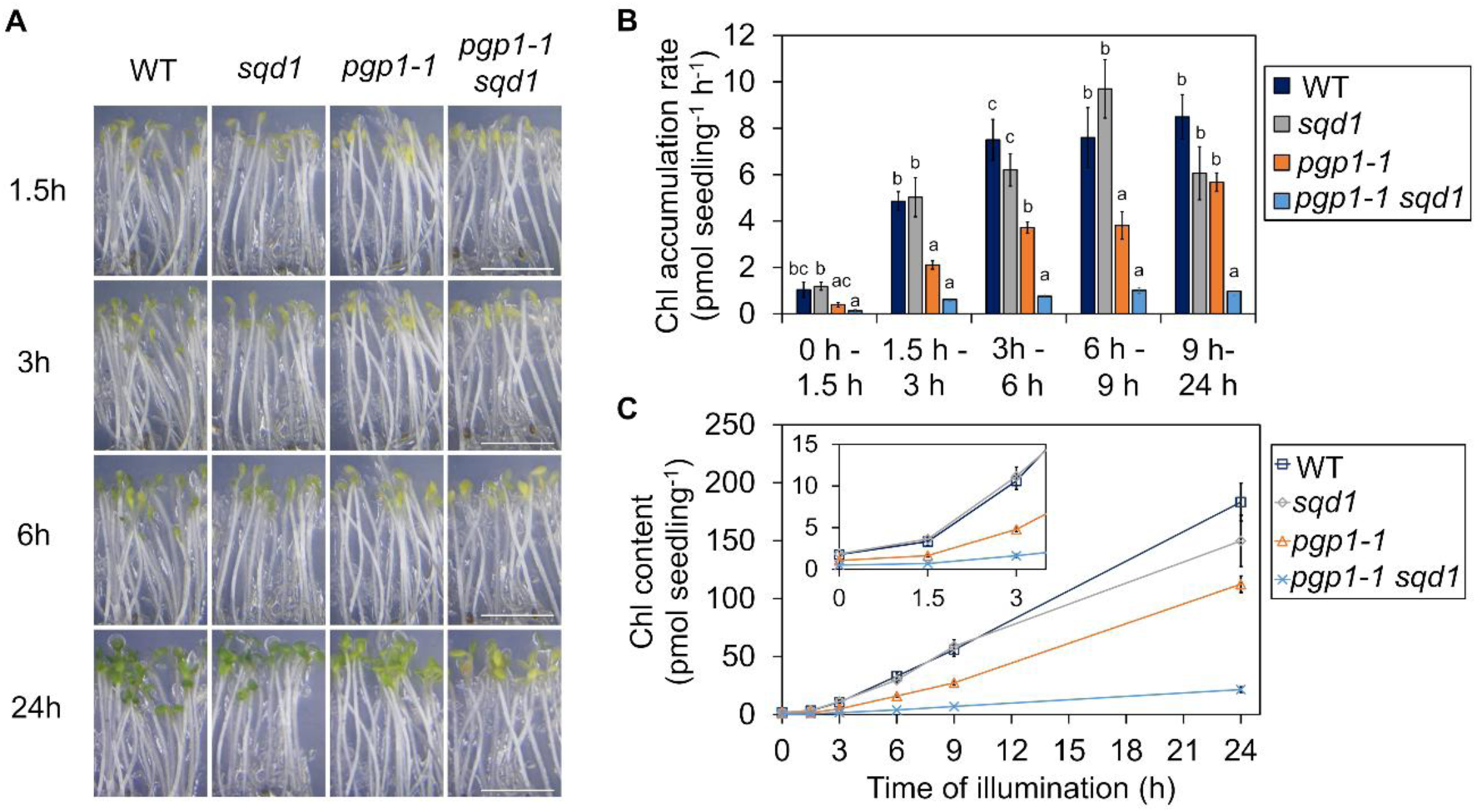
Chlorophyll (Chl) accumulation in 4-d-old etiolated cotyledons of wild-type (WT) and anionic lipid mutants during illumination. **(A)** Greening phenotype, **(B)** rate of Chl accumulation, and **(C)** Chl content in etiolated seedlings of WT, *sqd1*, *pgp1-1*, and *pgp1-1 sqd1* after 1.5, 3, 6, 9, and 24 h of continuous illumination. Chl content represents the sum of Chl *a* and Chl *b*. In **(C)**, the inset shows an enlarged view of the data at 0, 1.5, and 3 h. Protochlorophyllide content from Yoshihara *et al*. (2024) is shown for 0 h instead of Chl content. Data are means ± SE from 16 independent experiments at 0 h (Yoshihara *et al*., 2024) and 6 independent experiments at each time point from 1.5 to 24 h. Statistical significance was assessed using one-way ANOVA followed by Tukey’s honestly significant difference (HSD) test. In **(B)**, Different lowercase letters indicate significant differences among genotypes for each time period. For **(C)**, the results of the HSD tests are provided in Supplementary Table S3.

Next, we evaluated photosynthetic capacity during deetiolation using PAM Chl fluorescence measurements (Fig. 6). During the first 3 h of illumination, Chl fluorescence was too weak in all plants to yield reliable photosynthetic parameters. Between 6 and 24 h after illumination, *F_v_/F_m_* increased in *sqd1* and *pgp1-1* to levels similar to those in WT. Meanwhile, *pgp1-1 sqd1* seedlings consistently showed lower *F_v_/F_m_*than WT. Φ*_PSII_*, which represents PSII photochemical efficiency under actinic light and is affected by the overall performance of the photosynthetic electron transport system (Baker, 2008), was also lower in *pgp1-1 sqd1*. In contrast, *pgp1-1* showed higher Φ*_PSII_* than WT after 6 and 9 h of illumination, but no significant difference from WT at 24 h. In *pgp1-1* and *pgp1-1 sqd1*, Φ*_NPQ_*, which reflects regulated thermal dissipation of absorbed light energy, remained low throughout the illumination period, whereas in WT and *sqd1*, it was initially high and subsequently decreased to levels comparable to those in *pgp1-1* and *pgp1-1 sqd1* by 24 h. In *pgp1-1 sqd1*, Φ*_C_*, which reflects non-regulated energy dissipation, was substantially higher than in WT throughout the measurement, whereas Φ*_C_* in *pgp1-1* was only slightly higher during the first 9 h of illumination. These results suggest that, during the early stage of deetiolation, partial PG deficiency may even increase photochemical utilization of absorbed light energy along with decreased regulated energy dissipation, whereas additional loss of SQDG in the PG-deficient background severely perturbs the establishment of photosynthetic functions.

**Figure 6.**
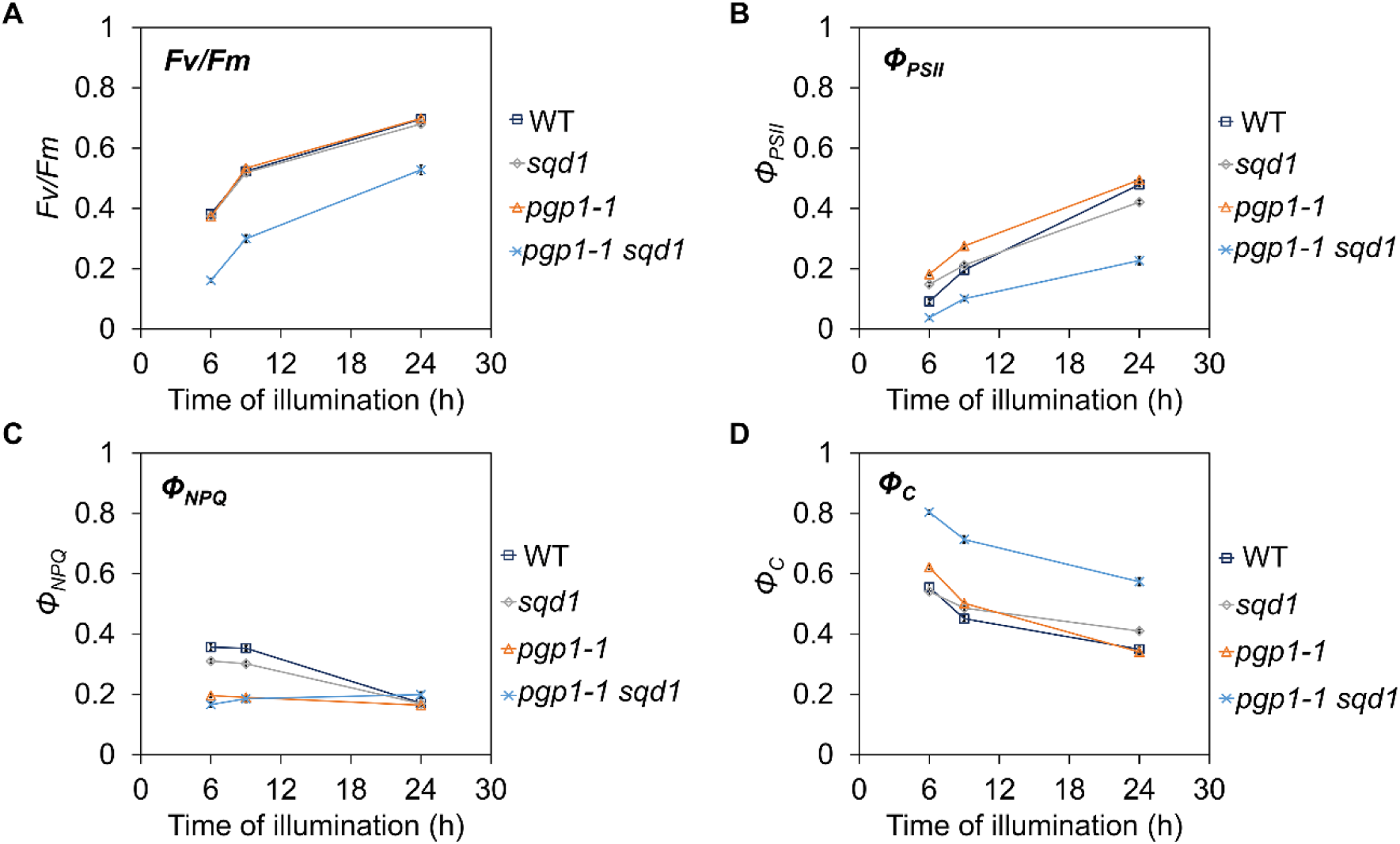
Establishment of the photosynthetic machinery in 4-d-old etiolated cotyledons of wild-type (WT) and anionic lipid mutants after illumination. **(A)** The maximum quantum yield of photosystem II (*F_v_*/*F_m_*), **(B)** the effective quantum yield of photosystem II (Φ*_PSII_*), **(C)** the quantum yield of regulated energy dissipation (Φ*_NPQ_*), and **(D)** the quantum yield of non-regulated energy dissipation (Φ*_c_*) in etiolated seedlings of WT, *sqd1*, *pgp1-1*, and *pgp1-1 sqd1* after 6, 9, and 24 h of continuous illumination. Data are means ± SE from approximately 50 individuals per genotype. Statistical significance was assessed using one-way ANOVA followed by Tukey’s honestly significant difference (HSD) test. The results of the post hoc tests are provided in Supplementary Table S4.

To investigate how the loss of plastid anionic lipid biosynthesis affects the behavior of Chl(ide)-protein complexes during deetiolation, we analyzed low-temperature fluorescence spectra of intact cotyledons after continuous illumination of etiolated seedlings (Fig. 7). WT and *sqd1* seedlings illuminated for 5 min showed a sharp fluorescence band peaking at 693 nm and a small shoulder around 680 nm, which are presumed to originate mainly from LPOR-bound and released Chl(ide), respectively (Lindsten *et al*., 1993; Gabruk *et al*., 2026). These plants exhibited a shift of the Chl(ide) peak to ∼685 nm within 20 min, followed by a gradual red shift to 689 nm during 6 h of illumination. In contrast, *pgp1-1* and *pgp1-1 sqd1* seedlings illuminated for 5 min showed a single fluorescence band peaking at 682 nm and 679 nm, respectively, consistent with the faster progression of the blue shift of Chl(ide) fluorescence after a single flash in these mutants (Fig. 2) (Yoshihara et al., 2024). After 10 min of continuous illumination, these mutants also exhibited a gradual red shift of the Chl(ide) peak, although *pgp1-1 sqd1* consistently showed shorter peak wavelengths than *pgp1-1*. Because PSII-LHCII complexes emit fluorescence at around 685-695 nm at 77 K (Lamb *et al*., 2018), the red shift of Chl fluorescence toward ∼690 nm after 1 h of illumination suggests the incorporation of Chl molecules into PSII-LHCII complexes. After 2 h of illumination, another fluorescence band appeared around 730 nm in all plants, which is known to originate from PSI-LHCI. At this time point, the peak wavelength of the PSI-LHCI band in *pgp1-1* and *pgp1-1 sqd1* was around 727 nm, shorter than those in WT and *sqd1* (∼731 nm). After 6 h of illumination, the PSI-LHCI band in *pgp1-1* and *pgp1-1 sqd1* shifted to ∼733 nm, similar to those in WT and *sqd1*, although the fluorescence intensity in *pgp1-1 sqd1* remained lower than in WT and *sqd1*. These results suggest that the organization and functional properties of PSI-LHCI complexes, including their interactions with PSII-LHCII, in *pgp1-1* and *pgp1-1 sqd1* differ from those in WT and *sqd1* during deetiolation.

**Figure 7.**
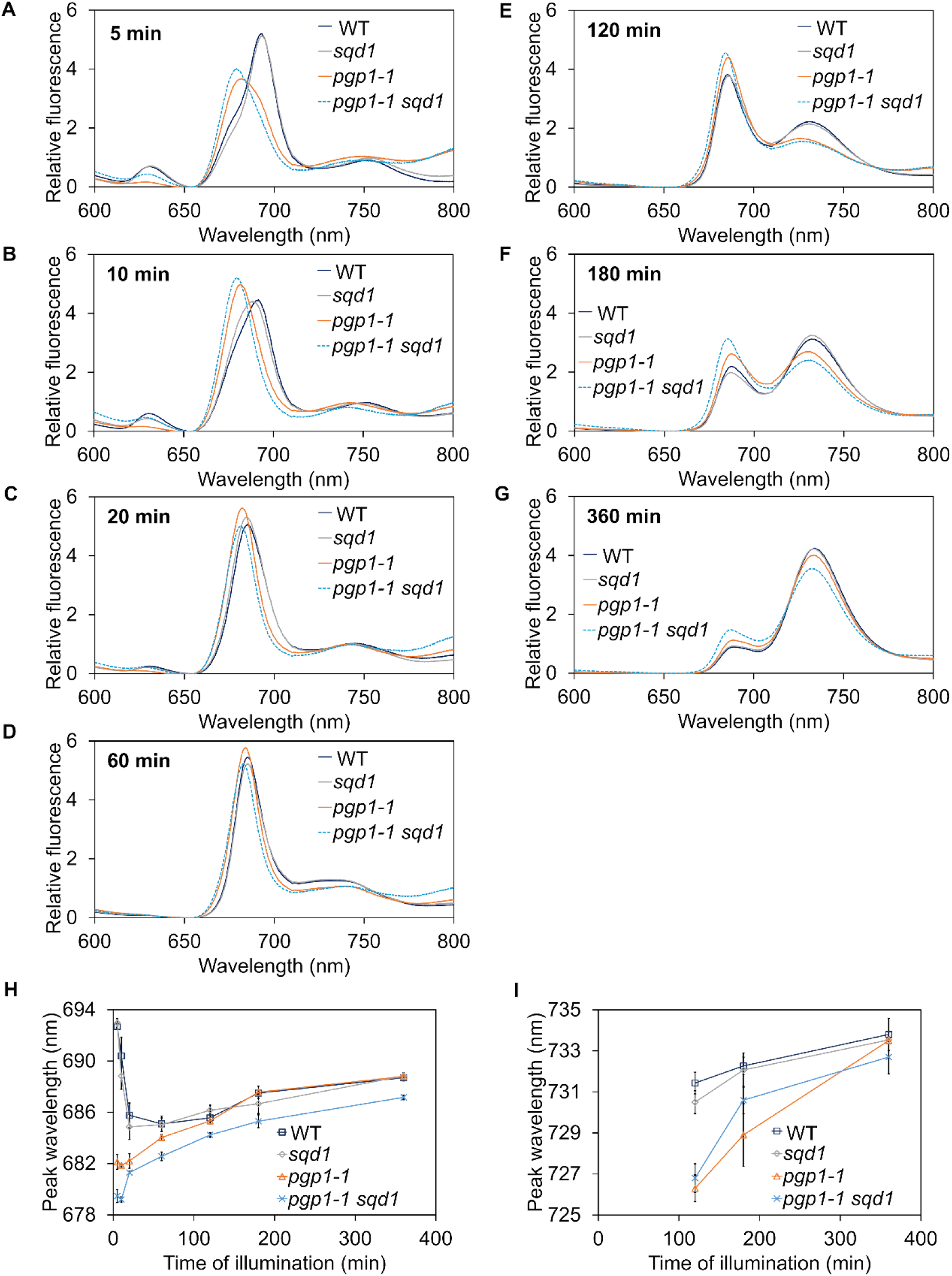
In situ low-temperature fluorescence spectra in 4-d-old etiolated seedlings of wild type (WT) and anionic lipid mutants after continuous illumination. 77 K fluorescence spectra of cotyledons after **(A)** 5, **(B)** 10, **(C)** 20, **(D)** 60, **(E)** 120, **(F)** 180, and **(G)** 360 min of continuous illumination. Fluorescence intensities were normalized to the mean fluorescence intensity over the 600–800 nm range. **(H, I)** Changes in fluorescence peak wavelength in the **(H)** 678–694 nm and **(I)** 725–735 nm regions during the illumination. Data are means ± SE from 3 independent experiments. Statistical significance was assessed using one-way ANOVA followed by Tukey’s honestly significant difference (HSD) test. The results of the post hoc tests are provided in Supplementary Table S5.

### SQDG is required for photosynthesis-associated gene expression when PG biosynthesis is reduced

Plastid lipid biosynthesis and subsequent membrane formation have been hypothesized to trigger photosynthesis-associated gene expression in both plastids and the nucleus (Fujii *et al*., 2025). To elucidate how decreased anionic lipid biosynthesis affects photosynthesis-associated genes encoded in plastids and the nucleus during deetiolation, we examined mRNA levels of key Chl biosynthesis genes *HEMA1* and *CHLH*, encoding the major isoform of glutamyl-tRNA reductase and the H subunit of Mg-chelatase, respectively; *LHCA4* and *LHCB6*, encoding subunits of the light-harvesting complexes of PSI and PSII, respectively; and *psaA* and *psbA*, encoding core subunits of the reaction centers of PSI and PSII, respectively (Fig. 8). *HEMA1*, *CHLH*, *LHCA4*, and *LHCB6* are encoded in the nucleus, whereas *psaA* and *psbA* are encoded in plastids. In WT, mRNA levels of all genes increased in response to illumination, and *sqd1* showed a similar upregulation pattern. In *pgp1-1*, the mRNA levels of some genes tended to be lower than those in WT at certain time points. Furthermore, *pgp1-1 sqd1* generally showed lower mRNA levels than WT across all examined genes during illumination.

**Figure 8.**
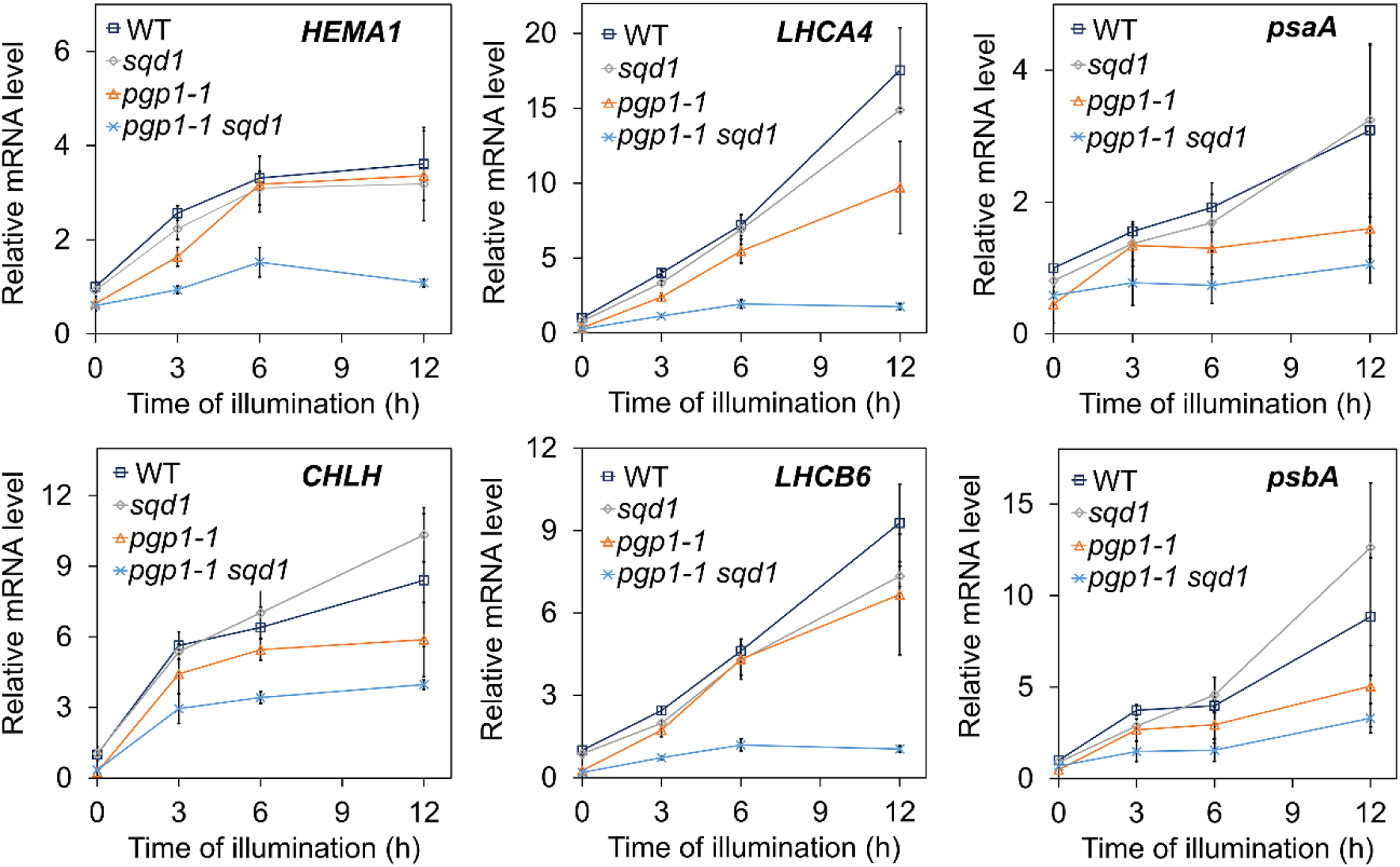
Induction of photosynthesis-associated gene expression in 4-d-old etiolated cotyledons of wild-type (WT) and anionic lipid mutants during illumination. The mRNA levels of photosynthesis-associated genes in etiolated seedlings before (0 h) and after (3, 6, 12 h) illumination are expressed as fold changes relative to the WT level at 0 h, after normalization to the reference genes *ACTIN8* and *UBIQUITIN11*. Data are means ±SE from 3 independent experiments. Statistical significance was assessed using one-way ANOVA followed by Tukey’s honestly significant difference (HSD) test. The results of the post hoc tests are provided in Supplementary Table S6.

## Discussion

In this study, we demonstrated that a partial reduction in plastid PG biosynthesis affects multiple processes during etioplast-to-chloroplast differentiation. Although etioplasts in *pgp1-1* seedlings were capable of forming apparently normal PLB lattice structures in darkness, these structures were highly susceptible to disassembly upon illumination, indicating reduced stability of the PLB architecture (Fig. 1A) (Yoshihara *et al*., 2024). Accordingly, the transformation from PLBs to lamellar membranes upon illumination (Fig. 3), along with the dissociation of Chlide-LPOR complexes and subsequent phytylation resulting in Chl (Fig. 2 and 4), proceeded more rapidly in *pgp1-1* than in WT. Furthermore, the more rapid rise in Φ*_PSII_* after the onset of illumination in *pgp1-1* than in WT may reflect a faster establishment of the photosynthetic electron transport machinery (Fig. 6B). However, impaired development of regulated energy dissipation during the early stage of deetiolation cannot be ruled out as a cause of the increased Φ*_PSII_* in *pgp1-1* (Fig. 6C). In contrast, Chl accumulation during illumination was delayed (Fig. 5), and the fluorescence properties of PSII-LHCII and PSI-LHCI, potentially reflecting changes in their development and interactions, were altered in *pgp1-1* (Fig. 7). Thus, although *pgp1-1* rapidly developed lamellar membranes from PLBs upon illumination, the development of the Chl biosynthetic pathway and photosynthetic apparatus differed markedly from that in WT. In addition, in *pgp1-1* cotyledon cells, a subset of plastids failed to develop thylakoid membranes and instead retained abnormal membrane structures during illumination (Fig. 1B and 3B). These findings suggest that PG is important not only for the structural integrity of PLBs but also for the coordinated progression of membrane remodeling and photosynthetic development during deetiolation.

The formation of regular PLB lattices despite their high susceptibility to light-induced disassembly indicates that PG is important for maintaining PLB structural stability rather than for their assembly. Upon illumination, Pchlide-LPOR-NADPH is instantaneously converted to Chlide-LPOR-NADP^+^, followed by rapid replacement of bound NADP^+^ by NADPH (Schoefs, 2001). The formation of Chlide-LPOR-NADPH complexes after photoreaction has been suggested to transiently stabilize oligomeric assemblies and delay PLB disassembly (Gabruk *et al*., 2026). Subsequently, these oligomeric complexes gradually dissociate, as reflected by the fluorescence peak shift from ∼695 nm to ∼680 nm at 77 K (the Shibata shift) (Shibata, 1957; Schoefs, 2001). In *pgp1-1* etioplasts, this fluorescence shift was accelerated (Fig. 2A) (Yoshihara *et al*., 2024), consistent with more rapid dissociation of the oligomeric Chlide-LPOR-NADPH complexes. Moreover, *pgp1-1* showed the rapid emergence of a fluorescence band around 675 nm, which presumably originates from free Chlide released from the complexes (Schoefs, 2001; Schoefs and Franck, 2008). Considering that PG increases the affinity between LPOR and NADPH in vitro (Gabruk *et al*., 2017), reduced PG biosynthesis in *pgp1-1* may destabilize the Chlide-LPOR-NADPH oligomers formed after illumination, thereby promoting rapid disassembly of the PLB lattice structure and release of free Chlide. Conversely, PG may contribute to delaying these processes by stabilizing oligomeric Chlide-LPOR-NADPH complexes.

The accelerated Shibata shift in *pgp1-1* is accompanied by faster Chlide-to-Chl conversion than in WT (Fig. 2D and Fig. 4B). In etioplasts, Chl synthase is present in both PLBs and prothylakoids, but the PLB-associated form is maintained in an inactive state (Lindsten *et al*., 1990). Because the addition of NADPH to isolated PLBs stabilizes oligomeric Chlide-LPOR-NADPH complexes and thereby inhibits the esterification of Chlide by Chl synthase, oligomerization of these complexes and the resulting PLB compaction are likely to interfere with Chl synthase activity (Lindsten *et al*., 1990, 1993). In *pgp1-1* etioplasts, the rapid dissociation of these oligomeric complexes and consequent disassembly of PLBs may allow

PLB-localized Chl synthase to become active more rapidly, resulting in faster Chlide-to-Chl conversion. In addition, the rapid release of free Chlide after photoconversion may contribute to its rapid phytylation by Chl synthase, while the alternative possibility that Chl synthase activation promotes Chl(ide) release cannot be excluded (Schoefs and Franck, 2008).

Despite the more rapid conversion of dark-accumulated Pchlide to Chl via Chlide in *pgp1-1*, de novo Chl biosynthesis after illumination was slower in this mutant than in WT (Fig. 5). Transcript levels of *HEMA1* and *CHLH*, which are representative genes tightly co-expressed with other key Chl biosynthesis genes (Kobayashi and Masuda, 2016), did not markedly decrease in *pgp1-1* (Fig. 8), implying that reduced expression of these genes does not account for the slower Chl accumulation. Meanwhile, we previously reported that Pchlide biosynthesis is retarded in *pgp1-1* etioplasts, particularly at the step of Mg insertion into protoporphyrin IX (Yoshihara *et al*., 2024). This impairment in the Chl biosynthetic pathway may persist during etioplast-to-chloroplast differentiation, resulting in reduced Chl accumulation rates despite accelerated activation of Chl synthase associated with PLB disassembly in *pgp1-1*.

In contrast to the slower Chl accumulation, Φ*_PSII_* in *pgp1-1* rose more rapidly than in WT during the first 9 h of illumination (Fig. 6B), whereas Φ*_NPQ_* was substantially lower (Fig. 6C). Because *F_v_*/*F_m_* increased similarly in WT and *pgp1-1* (Fig. 6A), the more rapid rise in Φ*_PSII_* in this mutant may reflect differences in the operation of the photosynthetic electron transport system rather than differences in the functional integrity of PSII. One possibility is that the photosynthetic electron transport components and/or associated metabolic processes, including ATP and NADPH synthesis and consumption, become functional more rapidly in *pgp1-1*, allowing more efficient electron transport and alleviating the conditions that trigger the development of non-photochemical quenching (NPQ) mechanisms. In this case, the rapid transformation of internal membranes in *pgp1-1* plastids during deetiolation might contribute to the more rapid establishment of these processes. Alternatively, mechanisms to induce NPQ development at the early stage of deetiolation may be impaired in *pgp1-1*, which could result in an increase in excitation energy input into PSII. If the photosynthetic electron transport system and downstream metabolic processes have sufficient capacity to accommodate this increased input, it could contribute to the higher Φ*_PSII_*, while the remaining excess energy would be dissipated through non-regulated pathways, consistent with the slightly higher Φ*_C_* in *pgp1-1* (Fig. 6D). In *Triticum aestivum*, energy-dependent NPQ and violaxanthin cycle operation increase during deetiolation, potentially helping to protect the developing photosynthetic apparatus from photooxidative damage (Garmash *et al*., 2013). A similar photoprotective role of NPQ may exist in *A. thaliana* during deetiolation, and its development could be impaired in *pgp1-1*. Further analyses are required to elucidate the molecular mechanisms underlying the altered partitioning of absorbed light energy in *pgp1-1* during the early stages of deetiolation.

Unlike *pgp1-1*, the *sqd1* mutant did not exhibit notable differences from the WT during etioplast-to-chloroplast differentiation, suggesting that SQDG alone is not essential for this process. However, the *pgp1-1 sqd1* double mutant showed more severe defects than *pgp1-1*. Many plastids in the double mutant failed to form regular PLB lattice structures even under infrared light fixation (Fig. 1B), and a substantial proportion retained abnormal membrane structures without developing thylakoid membranes after illumination (Fig. 3B). These observations suggest that a certain level of anionic lipids is required to maintain internal membrane integrity during plastid development and that SQDG plays a compensatory role under conditions of reduced PG biosynthesis.

Another prominent feature of *pgp1-1 sqd1* is its notably low *F_v_*/*F_m_*, indicating reduced PSII competence, which may also account for the low Φ*_PSII_* and high Φ*_C_* under light (Fig. 6). The shorter-wavelength peak of PSII-LHCII-derived fluorescence at 77 K after 6 h of illumination in *pgp1-1 sqd1* suggests altered energetic interactions between PSII and LHCII and/or within each component (Fig. 7, G and H). Structural studies of PSII-LHCII complexes have demonstrated that PG and SQDG molecules occupy multiple sites in PSII, including cavities around the plastoquinone binding sites, the monomer-monomer interface of the PSII dimer, and the interfaces between PSII and LHCII proteins (Wei *et al*., 2016; Su *et al*., 2017). In the thermophilic cyanobacterium *Thermosynechococcus elongatus*, the complete absence of SQDG causes partial impairment of PSII activity, with PG probably occupying all SQDG-binding sites (Nakajima *et al*., 2018), suggesting that PG can partially substitute for SQDG in maintaining PSII structure. Although it remains unclear to what extent SQDG can compensate for PG deficiency in the PSII-LHCII complexes, it is possible that PG and SQDG play partially complementary roles in preserving their structural integrity. If so, the complete absence of SQDG combined with partial PG deficiency in *pgp1-1 sqd1* may alter the lipid environment at anionic-lipid-binding sites in PSII-LHCII, thereby compromising the structural integrity of these complexes, as suggested by its low *F_v_*/*F_m_* (Fig. 6A). In addition, the attenuated light induction of *psaA* and *psbA*, and presumably of co-expressed genes encoding other photosystem proteins, might contribute to the impairment of photosynthetic establishment in *pgp1-1 sqd1* during deetiolation (Fig. 8).

The *pgp1-1 sqd1* mutant also exhibited substantially reduced Chl accumulation even compared with *pgp1-1* (Fig. 5). This reduction can be at least partly explained by a more severe defect in the Mg insertion step of the Chl biosynthetic pathway in *pgp1-1 sqd1* than in *pgp1-1* (Yoshihara *et al*., 2024). Although direct evidence is still lacking, the possibility that Chl biosynthesis is regulated through the thylakoid membrane as a scaffold has been discussed (Fujii *et al*., 2025). How changes in the lipid environment within plastids affect the functionalities of enzymes associated with the membrane-bound Chl biosynthetic pathway remains to be investigated. In addition, *pgp1-1 sqd1* showed strong attenuation of light-induced expression of *HEMA1* and *CHLH* (Fig. 8), which also likely contributes to the decreased Chl accumulation.

Previous studies using the knockout *pgp1-2* mutant indicate that PG biosynthesis is required for the light-induced expression of photosynthesis-associated plastid-encoded genes (PhAPGs) and nuclear-encoded genes (PhANGs) (Fujii *et al*., 2022). Moreover, decreased biosynthesis of either MGDG or DGDG attenuates the upregulation of PhAPGs and PhANGs during deetiolation, suggesting the importance of membrane lipids for these processes (Fujii *et al*., 2019*a*, 2024). The partial reduction in MGDG biosynthesis did not noticeably disrupt PLB formation in etioplasts or the initial PLB-to-lamellar membrane transformation during deetiolation (Fujii *et al*., 2017, 2019*a*), despite its negative effects on photosynthesis-associated gene expression. In contrast, overexpression of *CURVATURE THYLAKOID1A* disrupts PLB-to-thylakoid transformation without affecting PhAPG and PhANG expression (Sandoval-Ibáñez *et al*., 2021). These results suggest that the induction of PhAPG and PhANG expression during deetiolation is associated with membrane lipid biosynthesis but is not primarily dependent on the morphological alterations of pre-existing internal membranes. Considering that lipids are synthesized in the plastid envelope, de novo lipid biosynthesis at the envelope may contribute to triggering photosynthesis-associated gene expression at the onset of chloroplast development. Of note, loss of plastid PG biosynthesis in *pgp1-2* specifically disrupts the expression of PhAPGs that are transcribed by plastid-encoded RNA polymerase, including *psaA* and *psbA*, suggesting that PG biosynthesis is required for gene expression mediated by this polymerase (Fujii *et al*., 2022). Because the attenuation of *psaA* and *psbA* induction was more apparent in *pgp1-1 sqd1* than in *pgp1-1* (Fig. 8), SQDG may partially compensate for PG function in this process. In addition, plastid-to-nuclear retrograde signaling downstream of PhAPG expression has been implicated in the decreased PhANG expression observed in plastid lipid mutants (Fujii *et al*., 2022, 2024, 2025). Although different classes of plastid lipids play distinct roles in chloroplast development, all lipid classes are involved in upregulating photosynthesis-associated gene expression, suggesting that lipid biosynthesis itself, or the formation of an initial lipid bilayer resulting from this process, may trigger photosynthesis-associated gene expression during chloroplast development. Future studies will provide insights into the molecular mechanisms underlying this triggering process.

## Acknowledgments

We are grateful to our colleagues in the plastid dynamics research community (plastid.jp) for helpful and fruitful discussions.

## Author contributions

A.Y. performed most experiments, analyzed and interpreted data, and prepared the first draft of the manuscript. R. U., Ke.K., and N.N. performed transmission electron microscopic experiments. Ko.K. designed and conducted experiments, analyzed and interpreted data, and wrote and edited the manuscript.

## Data availability

The data underlying this article are available in the article and in its online supplementary material.

## Supplemental data

**Supplementary Video S1.** Sequential transmission electron microscopy images of etioplasts in a cotyledon of a 4-d-old dark-grown *pgp1-1 sqd1* seedling fixed under infrared light. The video comprises 63 images of consecutive ultrathin sections, each 80 nm thick.

**Supplementary Table S1.** Sequences of oligonucleotide primers used for quantitative reverse transcription PCR analysis.

**Supplementary Table S2.** Statistical analysis of the data shown in Fig. 2B and 2C.

**Supplementary Table S3.** Statistical analysis of the data shown in Fig. 5C.

**Supplementary Table S4.** Statistical analysis of the data shown in Fig. 6.

**Supplementary Table S5.** Statistical analysis of the data shown in Fig. 7H and 7I.

**Supplementary Table S6.** Statistical analysis of the data shown in Fig. 8.

